# Role of ethnicity and socio-economic status (SES) in the presentation of retinoblastoma: findings from the UK

**DOI:** 10.1101/625160

**Authors:** Rabia Bourkiza, Phillippa Cumberland, Hiranya Abeysekera, Manoj Parulekar, Mandeep S Sagoo, Jugnoo Rahi, M. Ashwin Reddy

## Abstract

**Purpose:** The aim of this study was to investigate if there was a relationship between ethnicity or socioeconomic status and the presentation of advanced non-familial retinoblastoma in the UK.

**Methods:** A cross sectional study based at the two centres providing retinoblastoma care in the UK. Non-familial cases of retinoblastoma (Rb) presenting between January 2006 and December 2011 were included. Data collected included: age at diagnosis, gender, child’s ethnicity, International Intraocular Retinoblastoma Classification (IIRC) stage with Groups D and E being considered advanced, laterality, treatment, and postcodes. Individual postcode (ZIP code) data was used to obtain the Index of Multiple Deprivation (IMD) score. A postal questionnaire was sent to participants’ parents to collect further, person-level, information on languages spoken and household socioeconomic position. Measures of severity of retinoblastoma also included: requirement for primary enucleation; the use of adjuvant chemotherapy; and mortality.

**Results:** 189 cases were analyzed. 98 (52%) male and 91 (48%) female. Median age at diagnosis was 16 months [IQR 8 – 34 months]. 153/189 (81%) of cases presented with advanced retinoblastoma; 75 (40%) group E, 78 (41%) group D. 134 (72%) of cases were treated with enucleation.

Multivariable analysis showed that older age at presentation was associated with enucleation and bilateral disease was associated with adjuvant chemotherapy. There was some indication that South Asian ethnicity and being in the most deprived IMD quintile were associated with a higher likelihood of presentation with advanced disease, but these estimates did not reach statistical significance.

**Conclusions:** In this first national UK study of patients with non-familial retinoblastoma, there was no evidence of an association of ethnicity or socio-economic status and the risk of presenting with advanced disease. This may reflect equality in access of health care in the UK. As a result, awareness campaigns should continue.

## Introduction

Retinoblastoma (Rb) is the most common primary intraocular malignancy of childhood worldwide,^1^ with approximately 50-60 new cases per year in the UK.

The International Intraocular Retinoblastoma Classification (IIRC) describes five groups of retinoblastoma (A to E),^2^ which represent the continuum of disease progression. Whilst globe salvage with focal treatments and/or a form of chemotherapy occurs in more than 90% for Groups A to C,^3^ the figure is just over 60% for group D eyes,^4^ and group E eyes (the most advanced form) are often enucleated at presentation. Retinoblastoma surgeons often elect to enucleate as 39% of patients with Group E eyes require adjuvant chemotherapy to reduce the risk of metastases.^5,6^ Thus, early diagnosis and prompt treatment is crucial for globe salvage, preservation of vision and reduced morbidity.

It is recognized that in resource poor countries increased lag time (time to diagnosis interval) is associated with increased mortality and extra-ocular Rb.^7,8^ Recently, it has been demonstrated in the UK that increased lag time for children with Rb is not associated with an increased risk of adjuvant chemotherapy post-enucleation nor higher frequency of Group E eyes.^9^

Similarly, low socioeconomic status has been stated as an important factor in the development of advanced disease in resource poor countries.^10^ In the United States, one study reported a trend for Hispanic children and children with no healthcare insurance to have more advanced disease although statistical significance was not achieved.^11^

In the UK, the National Health Service (NHS) aims to provide equal access to healthcare. We were keen to understand which ethnic or socioeconomic groups, if any, were presenting with advanced retinoblastoma leading to adverse outcomes including mortality, enucleation and adjuvant chemotherapy. Identification of specific groups would enable resources to be directed to these groups.

## Materials and Methods

### Study population and data collection

This was a national multicenter, retrospective, non-comparative study evaluating non-familial retinoblastoma cases in the United Kingdom (UK).

Two centres in London and Birmingham provide the National Retinoblastoma service to the UK population in which all affected children are treated. The registries at these two centres, the Royal London Hospital, Barts Health NHS Trust and Birmingham Children’s Hospital were reviewed and non-familial cases of Retinoblastoma presenting between 1st January 2006 and 31st December 2011 were identified. This allowed a minimum of 5-year follow-up to investigate mortality. Only non-familial cases were included in this study as screening is already available for first degree relatives of patients with retinoblastoma. As such presentation of familial retinoblastoma is not initiated by these families. The study was approved by the National Research Ethics Committee (Reference 11/LO/0981). This research adhered to the tenets of the Declaration of Helsinki. Written parental consent was obtained for inclusion of participants in the study.

Data collected on all patients from electronic patient records included: age at diagnosis (months), gender, child’s ethnicity, International Intraocular Retinoblastoma Classification (IIRC) stage at diagnosis ^2^, laterality, treatment, and postcodes. In cases of bilateral disease, the stage of the worse eye was recorded. Individual postcode (ZIP code) data was used to obtain the Index of Multiple Deprivation (IMD) score, a relative deprivation score based on residential location.^12^ Deprivation across 8414 geographical areas was assessed based on income, employment, health, education, housing, access and child poverty. Higher scores of IMD indicate higher socioeconomic status.

A postal questionnaire was also sent to participants’ parents to collect further, person-level, information on languages spoken at home and household socioeconomic position, including housing tenure, main carer’s educational qualifications and main wage-earner’s employment status and occupation coded using the Standard Occupational Classification from the UK Office of National Statistics.^13^

Parents were contacted twice by mail and at least twice by telephone if they did not respond to maximize completion and return of the questionnaire.

### Outcome measures

Measures of severity of retinoblastoma included: IIRC stage (A to E) at diagnosis with Groups D and E being considered advanced; requirement for primary enucleation; the use of adjuvant chemotherapy dependent upon presence of high-risk features for systemic spread on histopathological evaluation i.e. massive choroidal invasion (>3mm),^14^ retrolaminar optic nerve invasion or scleral invasion; and mortality.

### Statistical methods

Descriptive statistics are reported for the distribution of factors by severity of retinoblastoma, enucleation and adjuvant chemotherapy treatment. Multivariable logistic regression was used to model associations between the outcomes of interest and demographic and sociodemographic factors.

## Results

### Study sample characteristics

A total of 192 children with sporadic non-familial retinoblastoma presented in the UK over the six-year period of the study (1January 2006 to 31 December 2011). Three cases were excluded from analysis. Two were due to incomplete data as they emigrated, and one declined consent. No child died during the study period, and no child died within 5 years from diagnosis. Thus, 189 cases were available for the present study: 98 (52%) male and 91 (48%) female. (Table1) Median age at diagnosis was 16 months [IQR 8 – 34 months], range 1 month to 12 years and 2 months; 117 (62%) of cases presented in the first two years of life. There were 59 bilateral and 130 unilateral cases (67 right eye, and 63 left eye). Left eyes were the worst affected eye in bilateral cases, compared to right eyes (47% left eye vs 32% right eye); this difference was not statistically significant.

**Table 1:** Distribution of demographic and clinical factors for all cases (N=189), by severity and treatment.

|  | **Total**<br>**N = 189** |  | **Advanced IIRC**<br>**groups (D&E)**<br>**N = 153 (81%)** |  | **Enucleation**<br>**N = 134 (71%)** |  | **Adjuvant**<br>**chemotherapy**<br>**N = 68/134 (50%)** |  |
| --- | --- | --- | --- | --- | --- | --- | --- | --- |
|  | n | % | n | % | n | % | n | % |
| **Gender** |  |  |  |  |  |  |  |  |
| Female | 91 | 48 | 73 | 48 | 64 | 48 | 36 | 53 |
| Male | 98 | 52 | 80 | 52 | 70 | 52 | 32 | 47 |
| **Ethnicity** |  |  |  |  |  |  |  |  |
| English/Scottish/Welsh | 140 | 74 | 110 | 72 | 98 | 74 | 54 | 79 |
| Indian/Pakistani/<br>Bangladeshi | 23 | 12 | 21 | 14 | 18 | 14 | 9 | 13 |
| Black<br>Caribbean/African | 8 | 4 | 6 | 4 | 4 | 3 | 1 | 1 |
| Mixed White | 11 | 6 | 9 | 6 | 8 | 6 | 1 | 1 |
| Other White | 3 | 2 | 3 | 2 | 3 | 2 | 1 | 1 |
| Other | 3 | 2 | 3 | 2 | 2 | 1 | 2 | 3 |
| **IMD score (quintiles)** |  |  |  |  |  |  |  |  |
| 1 (most deprived) | 39 | 21 | 34 | 23 | 27 | 20 | 14 | 21 |
| 2 | 44 | 23 | 35 | 23 | 32 | 24 | 17 | 25 |
| 3 | 29 | 15 | 21 | 14 | 20 | 15 | 11 | 16 |
| 4 | 33 | 18 | 28 | 18 | 25 | 19 | 12 | 18 |
| 5 (least deprived) | 43 | 23 | 34 | 22 | 29 | 22 | 13 | 19 |
| **Age at diagnosis** |  |  |  |  |  |  |  |  |
| <1 yr | 71 | 37 | 52 | 34 | 41 | 31 | 27 | 40 |
| 1 to <2 years | 46 | 24 | 35 | 23 | 31 | 23 | 16 | 23 |
| 2 to <3 years | 32 | 17 | 29 | 19 | 28 | 21 | 10 | 15 |
| 3 to <4 years | 27 | 14 | 24 | 16 | 21 | 16 | 10 | 15 |
| 4 to 12 years | 13 | 6 | 13 | 8 | 13 | 9 | 5 | 7 |
| **Laterality** |  |  |  |  |  |  |  |  |
| Unilateral | 130 | 69 | 103 | 67 | 97 | 72 | 35 | 51 |
| Bilateral | 59 | 31 | 50 | 33 | 37 | 28 | 33 | 49 |

Overall, 153/189 (81%) of non-familial retinoblastoma cases presented with advanced retinoblastoma (IIRC groups D and E); 75 (40%) group E, 78 (41%) group D, 24 (13%) group C, 11 (6%) group B and 1 group A. The child with an IIRC A grade was detected by an optometrist on routine assessment. The majority, 134 (72%), of cases were treated with enucleation, 124 (93%) of whom had advanced disease. Those presenting with advanced retinoblastoma and those treated with enucleation were similarly distributed by demographic and socioeconomic factors to all cases (Table 1), as were those receiving adjuvant chemotherapy (68, 50% of those that were enucleated).

Multivariable analysis showed children 2 years or older and those with bilateral retinoblastoma were more likely to present with advanced disease (Table 2). Older age at presentation was associated with enucleation and bilateral disease with receipt of adjuvant chemotherapy. There was some indication that Indian/Pakistani/Bangladeshi ethnicity and being in the lowest (most deprived) IMD quintile were also associated with a higher likelihood of presentation with advanced disease, but these estimates did not reach statistical significance.

**Table 2:** Associations between advanced presentation retinoblastoma (grade D & E) and treatment, and demographic and socioeconomic factors. (N = 189)

|  | **Advanced IIRC grade**<br>**N = 153/189 (81%)** |  | **Enucleation \***<br>**N = 134/189 (71%)** |  | **Adjuvant chemotherapy \***<br>**N = 68/134 (50%)** |  |
| --- | --- | --- | --- | --- | --- | --- |
|  | **Unadj**<br>**OR** | **Adj OR** | **Unadj OR** | **Adj OR** | **Unadj OR** | **Adj OR** |
| **Gender** |  |  |  |  |  |  |
| Female | 1 |  | 1 | 1 | 1 | 1 |
| Male | 1.10 | 1.03 [0.46, 2.30] | 1.01 | 1.04 [0.47, 2.30] | 0.68 | 0.50 [0.22, 1.17] |
| **Ethnicity** |  |  |  |  |  |  |
| English/Scottish/Welsh | 1 | 1 | 1 | 1 | 1 | 1 |
| Indian/Pakistani/<br>Bangladeshi | 2.86 | 3.13 [0.62, 16] | 1.86 | 1.40 [0.36, 5.41] | 0.75 | 0.75 [0.22, 2.59] |
| Black<br>Caribbean/African | 0.82 | 0.59 [0.09, 3.72] | 0.41 | 0.16 [0.02, 1.18] | 0.28 | 0.35 [0.03, 4.20] |
| Mixed/Other White or<br>Other | 2.05 | 3.92 [0.77, 20] | 1.34 | 0.93 [0.23, 3.76] | 0.37 | 0.37 [0.09, 1.56] |
| **IMD score (quintiles)** |  |  |  |  |  |  |
| 1 (most deprived) | 1 | 1 | 1 | 1 | 1 | 1 |
| 2 | 0.57 | 0.61 [0.17, 2.15] | 1.19 | 1.90 [0.59, 6.17] | 1.13 | 0.94 [0.28, 3.14] |
| 3 | 0.39 | 0.35 [0.09, 1.37] | 1.11 | 2.55 [0.63, 10] | 1.22 | 1.03 [0.25, 4.28] |
| 4 | 0.82 | 0.79 [0.19, 3.39] | 1.59 | 1.99 [0.54, 7.37] | 0.92 | 0.55 [0.15, 2.04] |
| 5 (least deprived) | 0.55 | 0.57 [0.16, 2.07] | 0.99 | 1.19 [0.36, 3.94] | 0.81 | 0.66 [0.20, 2.25] |
| **Age at diagnosis** |  |  |  |  |  |  |
| <1 yr | 1 | 1 | 1 | 1 | 1 | 1 |
| 1 to <2 years | 1.16 | 1.58 [0.59, 4.25] | 1.62 | 1.39 [0.52, 3.74] | 0.52 | 0.66 [0.19, 2.25] |
| 2 to <3 years | 3.53 | **6.24 [1.50, 26]** | 5.12 | 3.58 [0.96, 13] | 0.29 | 0.66 [0.19, 2.30] |
| 3 to 12 years | 4.5 | **9.57 [2.34, 39]** | 6.22 | **4.30 [1.06, 17]** | 0.41 | 1.01 [0.31, 3.27] |
| **Laterality** |  |  |  |  |  |  |
| Unilateral | 1 | 1 | 1 | 1 | 1 | 1 |
| Bilateral | 1.46 | **3.24 [1.23, 8.57]** | 0.52 | 0.48 [0.19, 1.20] | 15 | **16 [4.8, 56]** |

Models adjusted for all factors in the table. \*Additional adjustment for IIRC groups

### Enhanced analysis of SES and Rb presentation (Questionnaire results)

The response rate was 42 % (79 responses out of 189). IIRC groups D or E were less likely to respond (39 % responded) compared to group A-C (53 % responded); Odds Ratio 0.57 [0.28, 1.20], which was not statistically significant.

Although there was a trend towards responding to the questionnaire with higher quantiles of IMD the results were not statistically significant (p = 0.06). With regards to ethnicity, compared to English/Scottish/Welsh, mixed race participants were more likely to respond (OR 6.36 [1.33, 30] p = 0.021). A summary of the questionnaire responses is presented in Table 3. There was no statistically significant association between language, employment status, social class, parental qualifications and accommodation tenure and outcomes (enucleation rate and adverse histopathology). Of note, there was no statistically significant association between the factors listed in Table 3 and advanced disease (IIRC Groups D and E).

**Table 3:** Distribution of household sociodemographic factors in subsample with enhanced individual level data on SES, by IIRC grade, enucleation and adjuvant chemotherapy. *Main wage earner ** State examination at age 16 *** State examination at age 18

|  | N | % D or E<br>IIRC grade | %<br>Enucleation | % Adjuvant<br>chemotherapy |
| --- | --- | --- | --- | --- |
| **Language spoken** |  |  |  |  |
| English only | 63 | 73.0 | 69.4 | 41.9 |
| English and other language | 15 | 86.7 | 66.7 | 36.4 |
| No English | 1 | 100 | 100 | 0 |
|  |  | p=0.46 | p=0.78 | p=0.67 |
| **Employment status\*** |  |  |  |  |
| Student/unemployed | 5 | 100 | 80.0 | 75 |
| Employed | 74 | 74.3 | 68.5 | 37.3 |
|  |  | p=0.94 | p=0.59 | p=0.14 |
| **Social class status\*** |  |  |  |  |
| Professional | 22 | 90.9 | 72.7 | 43.8 |
| Skilled | 28 | 60.7 | 64.3 | 42.1 |
| Semi-skilled | 11 | 81.8 | 72.7 | 25.0 |
| Manual | 9 | 77.8 | 75.0 | 33.0 |
|  |  | p=0.09 | p=0.89 | p=0.81 |
| **Highest qualification of main carer** |  |  |  |  |
| None | 8 | 87.5 | 75.0 | 33.3 |
| O Level** | 23 | 65.2 | 54.6 | 41.7 |
| A Level*** | 16 | 68.8 | 81.3 | 30.8 |
| Degree | 32 | 84.4 | 71.9 | 45.8 |
|  |  | p=0.29 | p=0.32 | p=0.82 |
| **Accommodation tenure** |  |  |  |  |
| Home owner | 59 | 76.3 | 65.5 | 46.2 |
| Rental accommodation | 20 | 75.0 | 80.0 | 25.0 |
|  |  | p=0.91 | p=0.23 | p=0.15 |

## Discussion

In this first national study of patients with non-familial retinoblastoma diagnosed over a six-year period to 2011 in the UK, there was no evidence of an association of ethnicity or socio-economic status and the risk of presenting with advanced disease.

A key strength of our study is that data was extracted from a prospective retinoblastoma database with no selection bias. In addition, we collected information on socioeconomic status directly from the patients’ parents to increase granularity of the data. For example, education is different from income and might help us with further interventions. Also, during this period of data collection enucleation rates were over 70% and we had information regarding high risk histopathological features. As globe salvage has increased due to new treatments (intra-arterial chemotherapy and intravitreal chemotherapy) such information is more difficult to acquire. However, the number of eyes that fall in the more advanced Groups D and E remain valid parameters to study.

In the US, it has recently been shown that there was a trend for Hispanics and those with unfavorable socioeconomic factors to have more advanced disease at presentation (more high risk adverse histopathology on local review). However, central review of histopathology slides did not provide evidence that this was statistically significant.^11^ This suggests that there may have been bias at local review particularly if the name of the child was not masked from the histopathologist. From 2000 to 2010, the data from 830 children were analyzed and an association between requirement for enucleation and being Hispanic and/or low SES existed.^15^ A retrospective analysis of the presentation of disease according IIRC (particularly groups D and E) may have been difficult to perform and the decision to enucleate was not standardized according to classification, thus bias on the part of the surgeon may have played a part. In addition, statistical significance was noted in mortality: 2 of 262 white children died (99% 5-year survival) compared to 6 of 89 black children (93% 5-year survival). The causes of mortality from retinoblastoma are again complex ranging from associated pinealoblastoma to treatment strategies and poor follow-up. Unfortunately, such details were not provided in that study.

Human Development Index for different countries correlates with survival for retinoblastoma.^16^ In Mexico, lower maternal education and poor prenatal housing conditions were significantly predictive of overall survival in unilateral disease, and more advanced IIRC grouping in bilateral disease, independent of diagnostic delay.^17^ In Brazil, maternal education carried a significant difference with outcomes (advanced stage at diagnosis, enucleation and survival).^18^ Interestingly, low SES *per se* was not associated with poorer outcomes.

We have previously shown that in the UK high-risk retinoblastoma (requiring adjuvant chemotherapy) is not associated with delayed lag time (time to diagnosis).^9^ This is counterintuitive but is found in all other paediatric cancers in high resource countries.^19^ Whereas low SES may be associated with more advanced disease at presentation in low resource countries or countries with unequal health care access, we wanted to understand if there were any vulnerable groups in a healthcare system that was free at point of access such as the UK. We have found no evidence of an association to suggest socioeconomic status or a certain ethnic group is disadvantaged. It is also difficult to argue that any particular ethnic group (present in the UK) is biologically more susceptible to advanced disease.

Our study draws on a national sample representative of the UK population of children with non-familial Rb. Nevertheless, power to detect true differences in risk of presentation with advanced disease may have been limited by the size of the sample. An inherent limitation to studying Rb is the rarity of the disease. As we hypothesized that ethnicity and socio-economic status may be risk factors, we undertook primary data collection on person/individual level SES factors, in order to allow deeper understanding of pathways. Unfortunately, we had only a moderate response to the questionnaire survey which limited our sample size further. There were some differences that failed to reach statistical significance. For example, there was some indication that Indian/Pakistani/Bangladeshi ethnicity, and being in the lowest IMD quintile were associated with higher likelihood of D/E IIRC grade, but this was not statistically significant. This may be due to the small study population size leading to inadequate power. However, the results are similar to the findings of another high resource country (USA) that has looked at this study question. ^11^

In summary, we report the largest cohort of patients with retinoblastoma in the UK with prospective data on ethnicity and socioeconomic status. Although, there is a trend between low SES and certain ethnic groups with advanced retinoblastoma, we have found no evidence of an association. This may reflect equality in access in primary health care. As a result, awareness campaigns highlighting the white reflex and strabismus should continue in their present format.

## Acknowledgments

Jugnoo Rahi is a National Institute for Health Research (NIHR) Senior Investigator. The views expressed in this article are those of the authors and not necessarily those of the NHS, the NIHR, or the Department of Health.

Research Grant from the Royal National Institute of the Blind, UK. The funding organization had no role in the design or conduct of this research. Phillippa M Cumberland is funded by the Ulverscroft Foundation

